# Novel function of TRIP6, in brain ciliogenesis

**DOI:** 10.1101/777052

**Authors:** Shalmali Shukla, Pavel Urbanek, Lucien Frappart, Ronny Hänold, Sigrun Nagel, Shamci Monajembashi, Paulius Grigaravicius, Woo Kee Min, Alicia Tapias, Olivier Kassel, Heike Heuer, Zhao-Qi Wang, Aspasia Ploubidou, Peter Herrlich

## Abstract

TRIP6, a member of the zyxin-family of LIM domain proteins, is a focal adhesion component. *trip6* deletion in the mouse revealed, unexpectedly, in view of its ubiquitous expression, a function in the brain: ependymal and choroid plexus epithelial cells were poorly developed, carrying fewer and shorter cilia, and the mice developed hydrocephalus. TRIP6 disruption, via RNAi or inhibition of its homodimerization, in a choroid plexus epithelial cell line, confirmed its function in ciliogenesis. Zyxin-family members carry numerous protein interaction domains. In common with assembly of other multiprotein complexes, ciliogenesis may be facilitated by molecular assembly factors. Super-resolution microscopy demonstrated TRIP6 localization at the pericentriolar material and along the ciliary axoneme. The requirement for homodimerization which doubles its interaction sites, its punctate localization along the axoneme, and its co-localization with other cilia components suggest a scaffold/co-transporter function for TRIP6 in cilia. This is the first discovery of a protein assembly factor essential for mammalian ciliogenesis.

## Introduction

Cell-Cell interactions, via adhesion between cells and between cells and extracellular matrix, are hallmarks of multicellular organisms, and are formed by multiprotein complexes at cellular membranes. Their nature is transient in that adhesions are disassembled and reformed in various processes such as cellular migration, cell division and tumorigenic transformation. The closeness of neighboring cells, also of different types, permits communication through gap junctions. Development and function of the brain are prime examples of such cell-cell interactions.

The formation of multiprotein complexes, including those involved in adhesions, often requires assembly factors. A family of scaffold proteins carrying several protein interaction domains serve as platforms for protein-protein interactions: the seven-member zyxin LIM domain family (including zyxin, lipoma-preferred partner LPP, Ajuba, WNT interacting protein WTIP, LIMD1, Migfilin and thyroid hormone receptor interacting protein [TRIP6]). Most of these proteins localize at focal adhesions and adherens junctions, compatible with their suspected role in the assembly of adhesion complexes. Their interaction partners and the cellular functions of these complexes, from actin organization/cell migration through tension sensing to signal transduction, have been elaborated in numerous cell culture experiments (e.g. Crawford et al., 1992; Schmeichel & Beckerle, 1994; Yi & Beckerle, 1998; Wang et al., 1999; Petit et al., 2000; Hirota et al., 2000; Yi et al., 2002; Guŕianova et al., 2005; Takizawa et al., 2006; Bai et al., 2007; Hirata et al., 2008; Lin & Lin, 2011; Dutta et al., 2018).

The zyxin family members show significant structural and functional similarities, and widely similar tissue specific expression pattern. Indeed, most of them seem to fully substitute for each other, as indicated by the viability and, by and large normal adult life, of mice with deletion of the genes for ZYXIN (Hoffman et al., 2003), LPP (Vervenne et al., 2009), Ajuba (Pratt et al., 2005), LIMD1 (Feng et al., 2007), and Migfilin (Moik et al., 2011).

In contrast to their similar expression in most tissues, the expression pattern of the zyxin family members in the brain differs: of all ZYXIN family members only *ajuba* and *trip6* were expressed in the embryonic and early postnatal brain showing a non-overlapping spatio-temporal pattern. *ajuba* RNA positive cells visualized by *in-situ* hybridization were mainly discovered in early embryogenesis (E10 to E13) and postnatally (mouse genome informatics atlas, Gray et al., 2004). By immunofluorescence microscopy TRIP6 was detected from E14.5 to early postnatal days with its expression restricted to the SOX2 positive cells of the ventricles (Lai et al., 2014). TRIP6 was reported to be transiently expressed in the embryonic hippocampus, the medial habenular nucleus, and the ventral posterior complex of the thalamus (Lv et al., 2015). In the adult mouse sensitive single cell RNA profiles from ventricular zone (VZ) identified ependymal cells with *trip6* expression, but also with *zyxin* and *ajuba* (Zywitza et al., 2018). Similar analysis for embryonic ventricular zone have not yet been done.

This embryonic brain-restricted and exclusive expression pattern between E14.5 and birth solely of TRIP6 among all ZYXIN family proteins, encouraged us to delete the gene in the mouse. While the mice were viable, similarly to the deletion of genes for the other zyxin family members, we observed a major phenotype: the mice developed an obstructive (non-communicating) hydrocephalus. Investigation into the underlying aetiology led us to the discovery of a novel and unexpected function of TRIP6 in brain ciliogenesis.

## Results

### TRIP6 expression in the brain

We confirmed *trip6* expression in the mouse brain by radioactive *in-situ*-hybridization (ISH) (**Figure 1A**). Widespread and intense *trip6* hybridization signals were found in the extracranial somatic tissues (see whole skull sections in **Figure S2A** and **B**). In the embryonic CNS (E14.5 to E18), however, *trip6* expression was highly restricted and limited to areas bordering the ventricles (**Figure 1A**, sagittal section, solid arrows), similar to published data (Lai et al., 2014). A similar *trip6* expression pattern persisted after birth (**Figure 1A,** P0-P6), albeit at reduced expression level, and was undetectable by *in situ* hybridization beyond P10 (not shown). *trip6* mRNA expression was also detected in choroid plexus, e.g. at E14.5, E18.5 and postnatal stages P0 and P3 (dashed arrows in **Figures 1A** and **S2A** and **B**). Immunostaining of the protein confirmed the *in-situ*-hybridization data and identified the exact region and cell type expressing TRIP6: a thin layer of cells bordering the ventricles in the ventricular zone (VZ) and in the choroid plexus anlage, both during embryogenesis (**Figure S2C**) and postnatally (**Figure 1B**).

**Figure 1:**
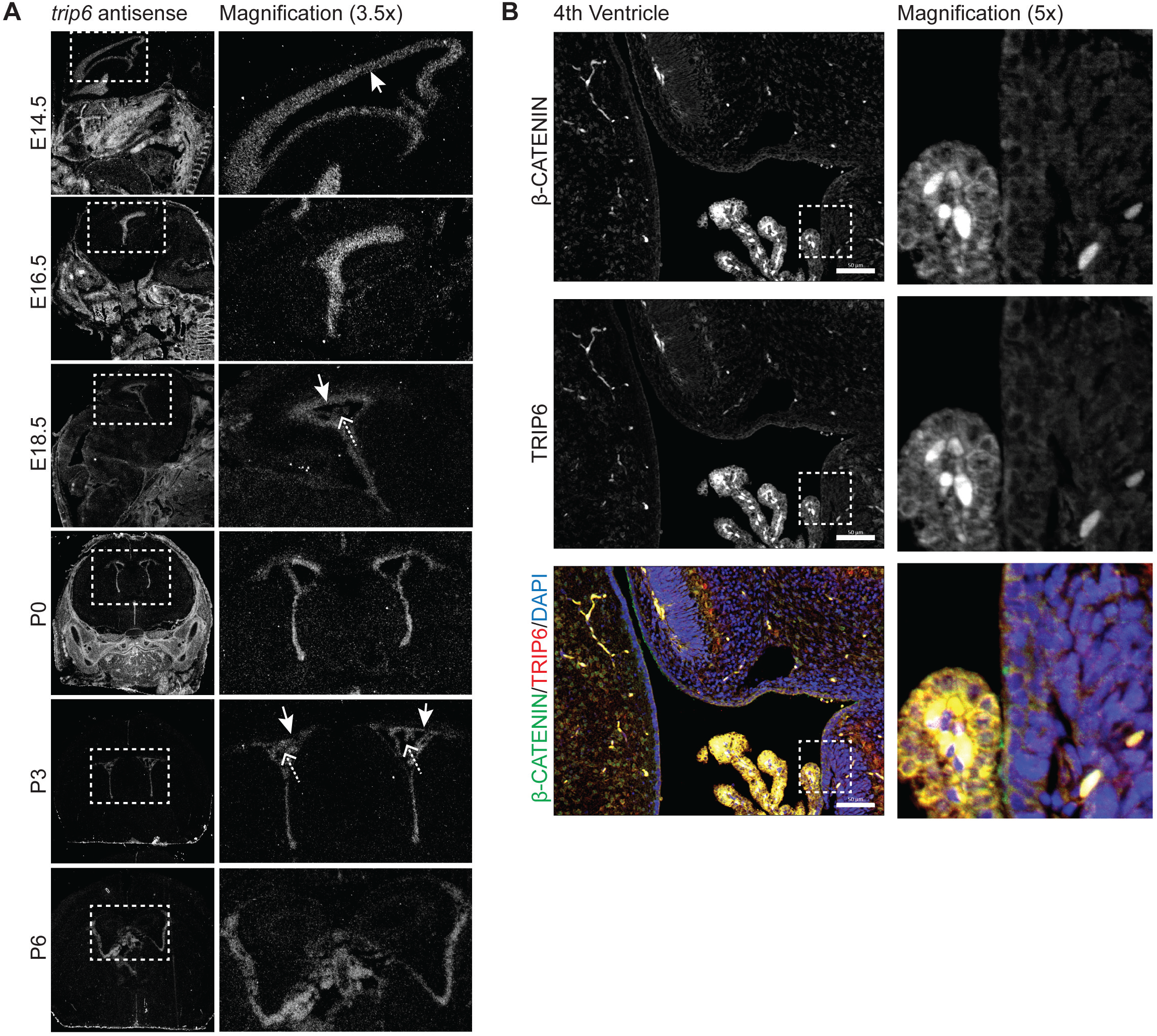
Expression of *trip6* in embryonic and postnatal brain. (**A**) mRNA expression in mouse brain sections by *in situ* hybridization (ISH): embryo sagittal sections E14.5 to E18.5, postnatal coronal sections P0 to P6. Arrows point at ventricular and subventricular zone (VZ/SVZ), dashed arrows point at choroid plexus. Control sense ISH is shown in Figure S2A and B. (**B**) TRIP6 protein expression determined by anti-TRIP6 antibody and co-localization of β-catenin in sagittal sections through the 4th ventricle at P4. Nuclei were stained with DAPI. The encircled areas are shown in magnification. Scale bar: 50μm (B).

TRIP6 co-localized with β-catenin (**Figure 1B, S2C**) and N-cadherin (**Figure S2C**), marking predominantly the cellular membranes of both ependymal cells and choroid plexus epithelium. Oblique sections tangentially through the VZ displayed the net-like structure of VZ cell membrane staining for β-catenin, N-cadherin and TRIP6 (**Figure S2C**). Thus, TRIP6 is expressed in cellular adhesion complexes along the ventricle and in the choroid plexus epithelium.

### *trip6* gene deletion in the mouse results in formation of hydrocephalus

The generation of *trip6*^−/−^ mice by standard technology is outlined in **Figure S1**. The *trip6*^−/−^ mice were viable through birth. While heterozygous animals were indistinguishable from their wild type littermates, a large fraction of *trip6* knock-out mice developed a drastic phenotype: hydrocephalus (**Figure 2**). In the C57BL/6 background, the hydrocephalus incidence was 43 % (**Figure 2A,B**). Haematoxilin-Eosin (H&E) stained brain tissue revealed the enormous extension of the lateral ventricle in these animals (**Figure 2C**). The overall brain morphology was not altered in the fraction of mice escaping the formation of hydrocephalus (not shown).

**Figure 2:**
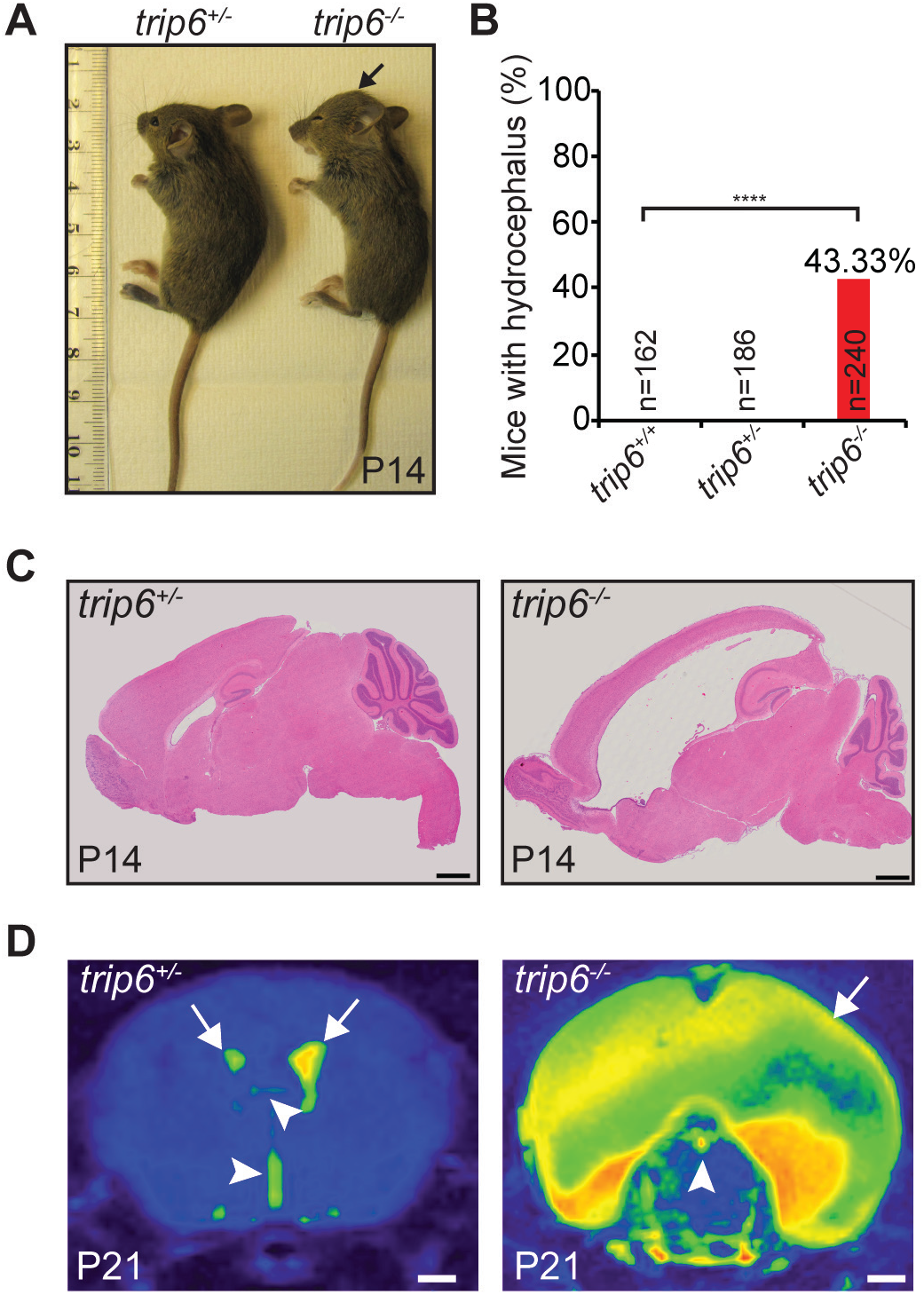
*trip6* deletion causes perinatal hydrocephalus. (**A**) Postnatally the bulging of the skull becomes visible. (**B**) No spontaneous hydrocephalus was detected in wild type and heterozygote mice. Not all mice with *trip6* deletion suffered from hydrocephalus. The frequency in *trip6*^−/−^mice was 43.33 %. n = number of mice, (p<0.0001 chi squared test). (**C**) H&E stained sagittal sections of P14 brains show enlarged lateral ventricles in *trip6*^−/−^compared to *trip6*^+/−^mouse. (**D**) Axial brain MRI images from a P21 control mouse and a P21 hydrocephalus carrier at the level of the lateral ventricles (arrow) and third ventricle (arrowhead depicting enlargement of the lateral ventricles in *trip6*^−/−^mice. Scale bar: 1mm (C and D).

High resolution MRI showed that the lateral ventricles were enormously enlarged, whereas in the third and fourth ventricles no increased fluid accumulation was detected (**Figure 2D**). Maximal ventricle enlargement was reached at P24 (data not shown). This phenotype speaks for a CSF circulation block in the foramen of Monro (upstream of the third ventricle) and/or in the aqueduct of Sylvius, between third and fourth ventricle (reviewed by Kousi & Katsanis, 2016) - suggesting the existence of an obstructive hydrocephalus.

As congenital hydrocephalus is often accompanied by delamination of ependymal cells from the dorsal wall of the aqueduct with fusion of the ventral and dorsal walls and consequently aqueduct obliteration (Wagner et al., 2013; Páez et al., 2007), we focused on analysis of the ependymal cell layers.

### Defective ependymal cells in *trip6*^−/−^ brains

We examined the ependyma and choroid plexus of early postnatal stages. Ependymal cells differentiate late in embryogenesis, e.g. between E18.5 and birth (reviewed by Narita & Takeda, 2015). Both terminally differentiated ependymal cells and choroid plexus epithelial cells carry motile cilia. H&E staining of *trip6*^−/−^ brain sections at P14 showed a considerably thinner and more flat layer of ependymal cells as compared to *trip6*^+/−^ control (**Figure 3A**). The number of cilia appeared to be reduced, and those cilia present were shorter. Immunofluorescence microscopy of S100β and acetylated α -tubulin (Ac. TUB), ependymal and cilia marker respectively (**Figure 3C-G**) confirmed the alterations in ependymal layer and the cilia defects (**Figure 3F, G**, compare with *trip6*^+/−^ in **E**). At these magnifications by which individual cilia cannot be resolved, we observed clusters of cilia at the *trip6*^+/−^ ependyme (**Figure 3E**) that were reduced in *trip6*^−/−^(**Figure 3F, G**).

**Figure 3:**
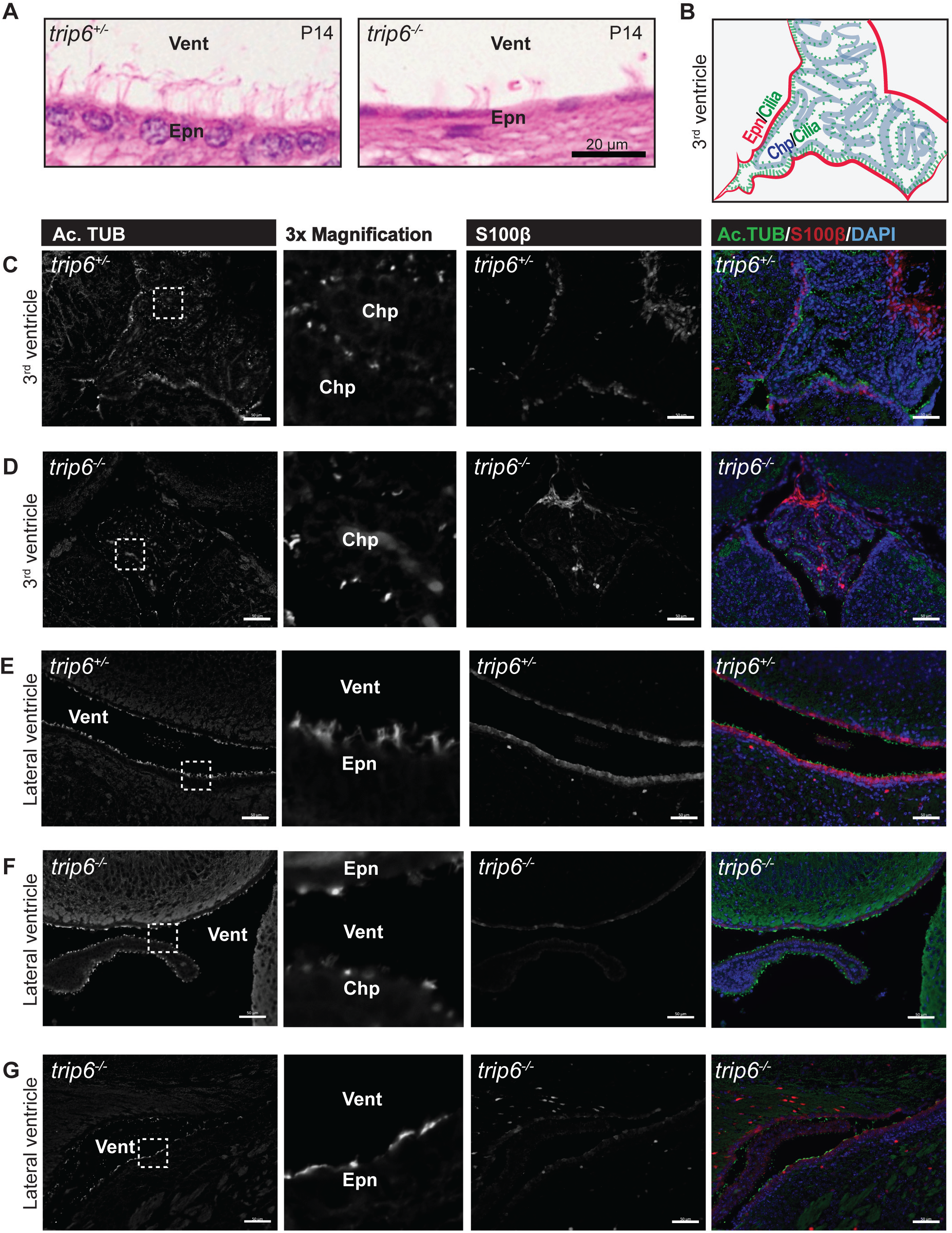
Defective differentiation of ependyma and choroid plexus in *trip6*^−/−^mice. (**A**) H&E stained sections show reduced cell size and fewer number of cilia in *trip6*^−/−^ependyma. (**B**) cartoon for the section shown in C. (**C**) and (**D**) Representative images of 3rd ventricle of *trip6*^+/−^ (**C**) and *trip6*^−/−^(**D**) P4 mice. *trip6* deletion lead to reduction of cilia (Ac.TUB) and diminished S100β positive ependymal cells. Representative images of lateral ventricles of *trip6*^+/−^(**E**), *trip6*^−/−^ hydrocephalic mouse (**F**) *trip6*^−/−^mouse without hydrocephalus. Note the reduced cilia (Ac. TUB) and ependymla cells (S100β) cells in the mutant mice. For the expression of Acetyl-tubulin magnified areas are shown. Vent = ventricle; Epn = ependyme; Chp = choroid plexus; Ac.TUB = acetylated α-tubulin. Scale bars: 20μm (A), 50μm (C to G).

Could the ependymal layer thinning and cilia reduction be simply the result of enhanced intracranial pressure in the lateral ventricles caused by the hydrocephalic volume increase? This hypothesis, however, could be ruled out because a similar thinning of ependyma was also seen in brains of *trip6* knock-out mice that did not develop enlarged lateral ventricles (**Figure 3G**; compare with section from mouse with hydrocephalus in **Figure 3F**), as well as in the third ventricle of *trip6*^−/−^mice downstream of the circulation block where there is no increase in the intracranial pressure (compare *trip*^+/−^in **Figure 3C** with *trip6*^−/−^in **D**). The ependyme (S100β) and its cilia (Ac. TUB) were observed to be reduced in *trip6*^−/−^animals (**Figure 3D**) compared to *trip6*^+/−^mice (**Figure 3C**, to help with the orientation, a cartoon is shown in **Figure 3B**). Deletion of *trip6* reduced also the cilia at the choroid plexus epithelial cells (compare **Figure 3C** and **D**). From screening through the VZ of several sections (H&E and IF staining) it appears that the ependymal layer of the *trip6*^−/−^brains showed patches of interruption as if the cells were sometimes dissociated from the basement membrane (not shown). Altogether, these data suggest that the differentiation of both ependyma and choroid plexus are negatively affected by the deletion of *trip6*.

### *trip6* deletion does not affect adherens junctions

In wild type mice TRIP6 co-localized with β-catenin and N-cadherin (**Figures 1B and S2C**). This was to be expected given the role of TRIP6 in the assembly of adhesion complexes in cell culture (Guryanova et al., 2005; Bai et al., 2007). The role of adhesion complexes in ciliogenesis is well documented (Antoniades et al., 2014). Thus, in view of the ciliary malformation of choroid plexus and ependymal cells in *trip6*^−/−^mice, we asked whether the adhesions were affected? Surprisingly, localization of β-catenin at cellular membranes was not altered in *trip6*^−/−^mice (**Figure 4**, left column) indicating that cell-to-cell adhesion does not - at least not structurally - depend on TRIP6.

### TRIP6 is associated with ciliary structures

Clusterin-associated protein 1 (CLUAP1, also called IFT38) is a component of the intraflagellar transport complex B (IFTB) which is involved in transfer of components into and out of the ciliary axoneme (Botilde et al., 2013; Katoh et al., 2016). CLUAP1 marked the apical rim of the ependymal layer carrying the motile cilia in the *trip6*^+/−^brain sections (**Figure 4**, third columns) resembling the staining of acetylated-tubulin (**Figure 3**). In sections from *trip6*^−/−^mice CLUAP1 staining was reduced (**Figure 4**), compatible with the reduction of acetylated tubulin labelled cilia.

**Figure 4:**
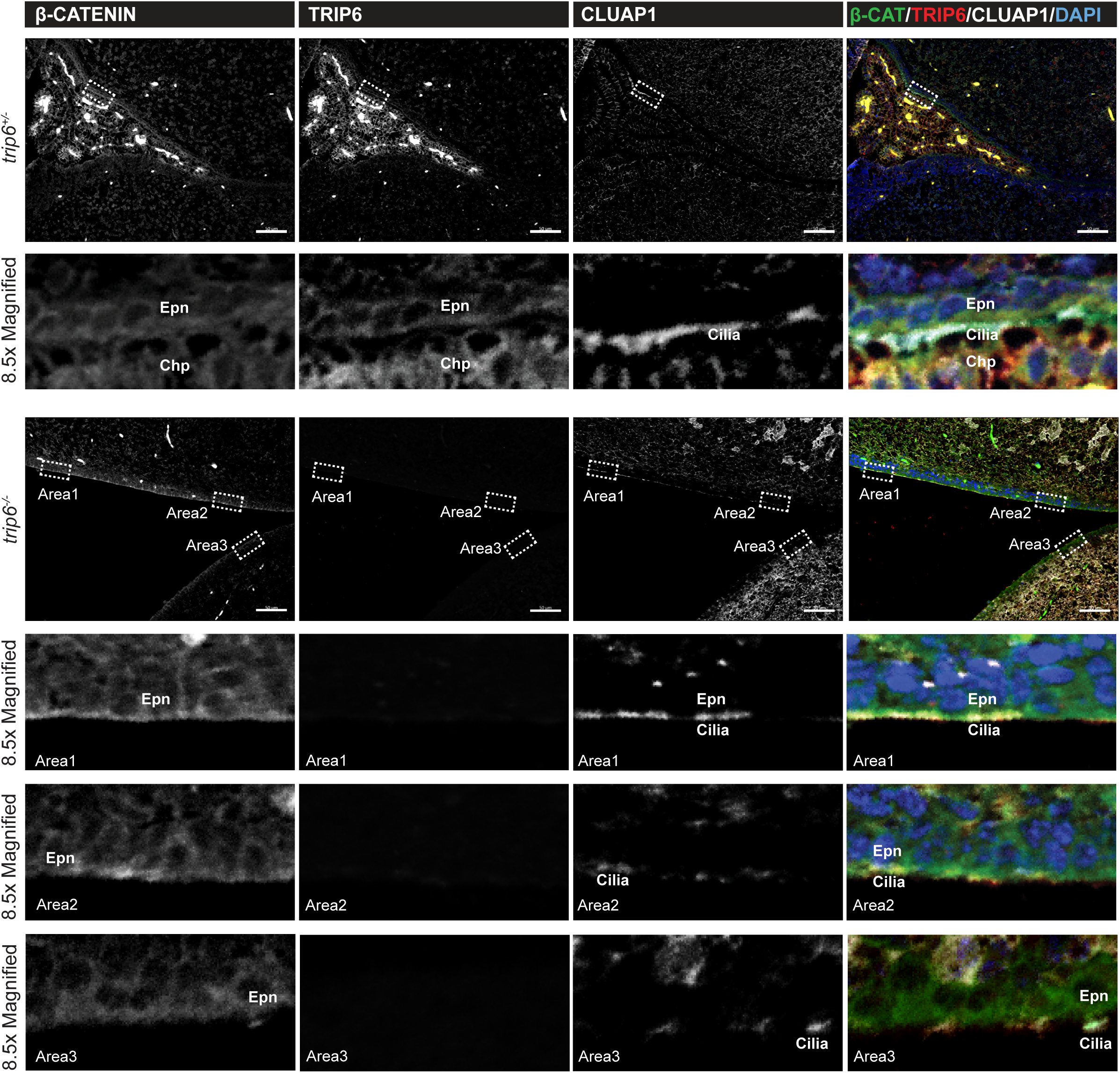
*trip6* deletion reduces ciliogenesis but does not affect cell-cell adhesion. Lateral ventricle sections of *trip6*^+/−^and *trip6*^−/−^mice (P4) were analyzed for expression of β-catenin, TRIP6 and the ciliary protein CLUAP1. Selected areas were magnified and oriented such that the ventricular lumen was towards the bottom. While in *trip6*^+/−^CLUAP1 marks a fairly continuous cluster of cilia, CLUAP1+ cilia were reduced and patchwise lacking in *trip6*^−/−^ventricles. Epn = ependyme; Chp = choroid plexus. Scale bar: 50μm.

So far we have documented that choroid plexus and ependymal cells were mal-formed in the absence of TRIP6 and that they lacked the normal abundance of motile cilia. At which level does TRIP6 act?

The decisive observation was obtained by co-immunofluorescence microscopy: acetyl-tubulin (**Figure 5A)**, CLUAP1 (**Figure 5B)** and TRIP6 co-stained cilia clusters (**Figure 5A** and **B**). In brain sections from *trip6*^+/−^mice TRIP6 localized in the cilia stained for acetylated tubulin as well as CLUAP1 (**Figure 5A,B**). These data suggest that TRIP6 is part of the ciliary structure. A more accurate localization of TRIP6 was obtained by super-resolution microscopy. “Dots” of TRIP6 were detected at the base, along the length and at the tip of the cilium of lateral ventricle ependymal cells (**Figure 5C,D**). The punctate pattern of the TRIP6 staining differed from one cilium to the other possibly capturing different locations of TRIP6 in the process of intraflagellar transport. In *trip6*^−/−^mice, however, acetylated tubulin stained cilia clusters of the ependyme were reduced (**Figure 5D**). The acetylated tubulin accumulated along the ventricular surface of the ependymal cells, but the thin layer suggests that the ciliary axoneme failed to elongate (**Figure 5D**).

**Figure 5:**
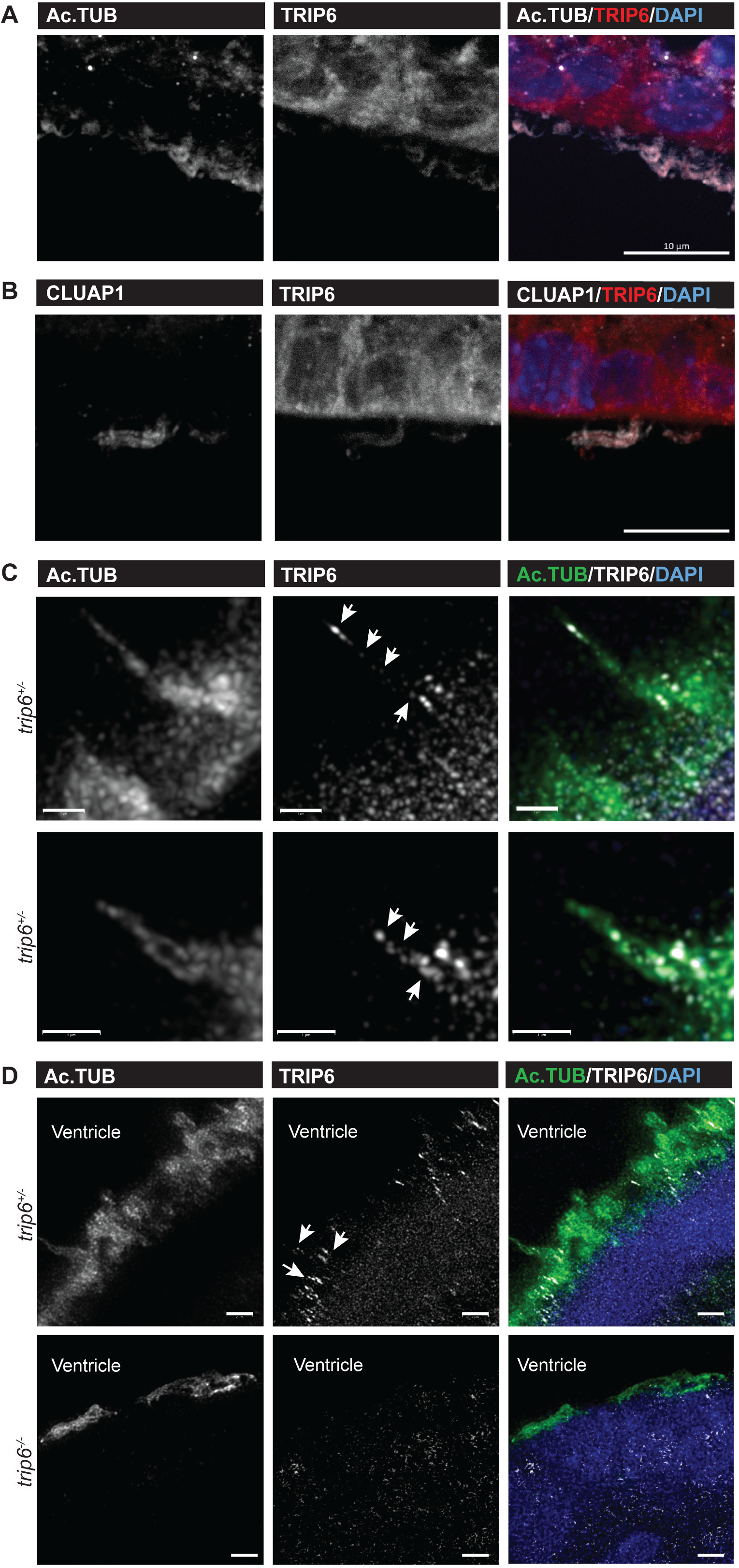
Association of TRIP6 with ciliary structure. Immunofluorescence for acetyl-tubulin (Ac.TUB) (**A**) CLUAP1 (**B**), and TRIP6 show co-localization in cilia clusters. (**C**) Super resolution imaging of cilia of individual cilia shows that Ac.TUB and TRIP6 co-localize in cilia. While Ac.TUB stained more uniformly, Trip6 was found at different locations along individual cilia. (**D**) Super resolution imaging of ependymal cilia from *trip6*^+/−^ and *trip6*^−/−^ mice (P4). Ac. TUB was accumulated along the ventricular surface of the cell but the axoneme failed to elongate in *trip6*^−/−^ mice. Scale bars: 10μm (A and B), 1μm (C) and 2μm (D).

### TRIP6 functions in ciliogenesis

To further investigate the putative roles of TRIP6 in ciliogenesis, we used the murine choroid plexus cell line Z310 (Zheng & Zhao, 2002). Its monolayer culture presented similar β-catenin staining (**Figure 6A**) as seen in choroid plexus epithelial cells and ependymal cells of *trip6*^+/−^mice (**Figures 1** and **S2**) with the cadherin-β-catenin complexes (assayed via β-catenin staining) indicating cell-cell adhesion. In agreement with published reports (Takizawa et al., 2006, Bai et al., 2007), TRIP6 also localized at focal adhesions as illustrated by anti-paxillin immunofluorescence staining (**Figure 6B** and **S3B**).

**Figure 6:**
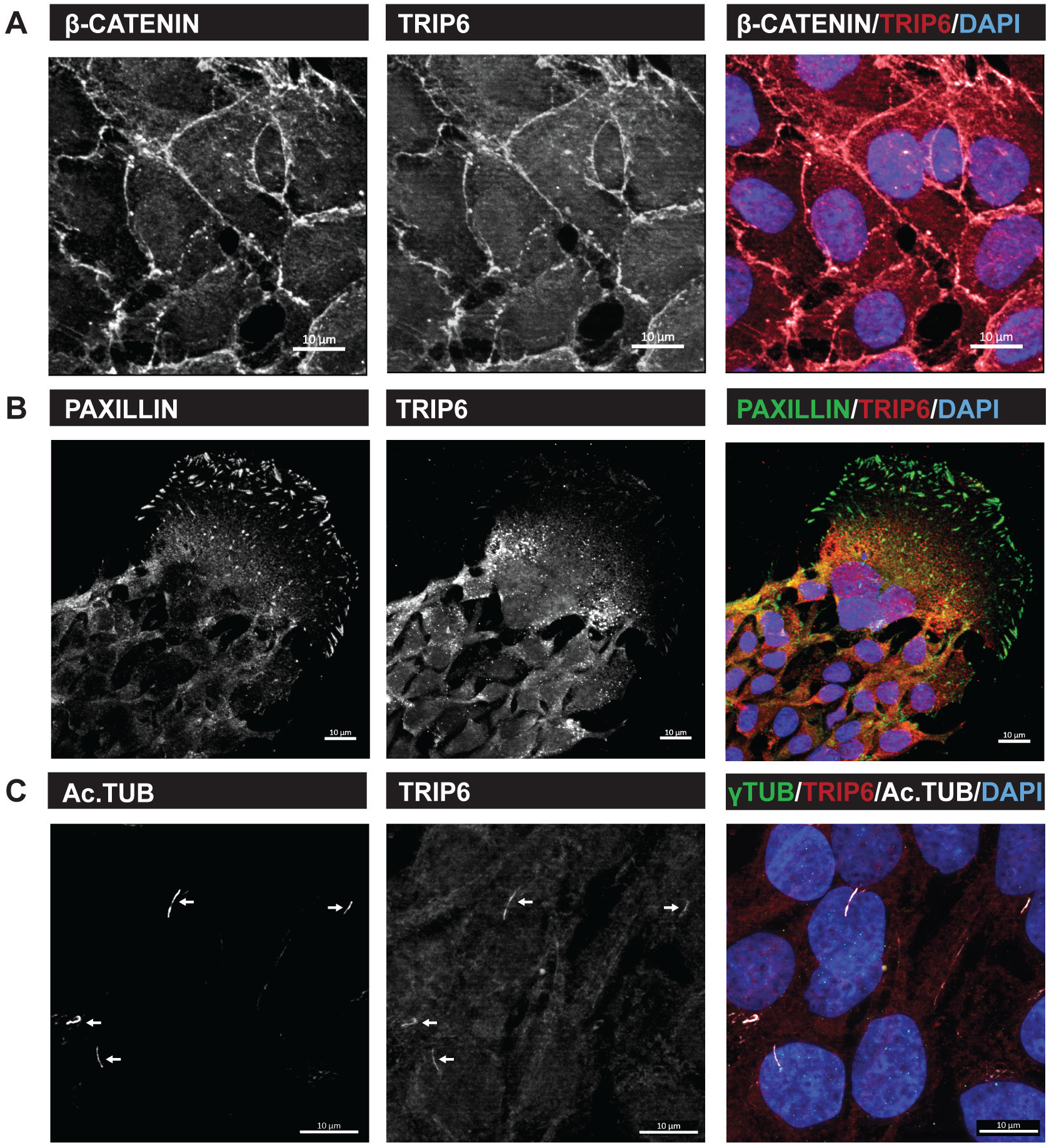
TRIP6 expression in adhesion complexes and cilia in Z310 cells. (**A**) Z310 cells were cultivated as described in Material & Methods. β-catenin and TRIP6 co-localize at adherens junctions. (**B**) Paxillin and TRIP6 are found at focal adhesions. (**C**) Induced cilium formation by starvation of Z310 cells. Cilia were identified by the basal body marker γ-tubulin and the axoneme marker acetyl-tubulin (Ac.TUB). TRIP6 co-localizes with Ac. TUB. Scale bars: 10μm (A, B and C).

Upon serum starvation, Z310 cells developed primary cilia **(Figure 6C**) that showed similar acetylated tubulin and TRIP6 localization (marked by arrows) as found in choroid plexus epithelial cells in brain sections (**Figure 5**). Is TRIP6 required for the formation of cilia formation in these cultured cells? Treatment of Z310 cells with *trip6* siRNA (siTrip6) resulted in cilia of reduced number and length compared to cells transfected with non-targeted siRNA (**Figure 7A**, compare siTrip6 vs siControl). Most cells transfected with *trip6*-specific siRNA produced only “puncta” which consisted of basal bodies with unextended axoneme marked by acetyl-tubulin/ARL13B (the cilium component ADP ribosylation factor like GTPase 13B; Roy et al., 2017). Quantification (**Figure 7A**) showed that *trip6* siRNA reduced the number of cells carrying cilia (left panel). The fraction of cells with elongated cilia was diminished (middle panel) and - in those cells that formed cilia - the cilium length was very significantly reduced (**Figure 7A**, right panel). Thus, TRIP6 is required for cilium formation and elongation.

**Figure 7:**
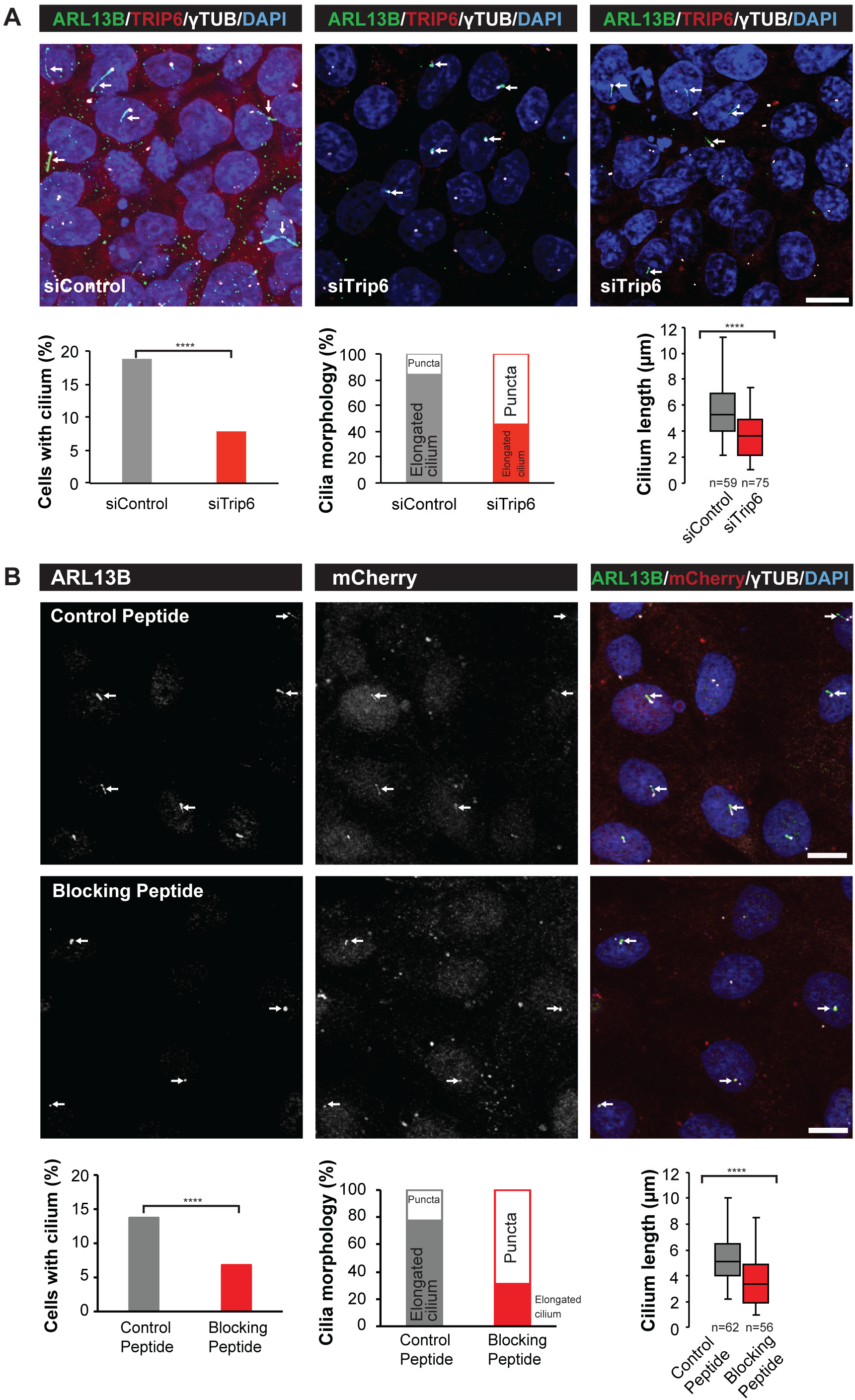
*trip6* downregulation or blocking TRIP6 dimerization in Z310 cells result in defective ciliogenesis and cilium elongation. (**A**) Cilia were induced by starvation in cells either treated with control siRNA or *trip6* siRNA. Cilia were identified by antibodies against γ-tubulin (basal body) and ARL13B (axoneme). Anti-TRIP6 was used to visualize the downregulation of TRIP6. γ-tubulin/ARL13B+ Cilia were counted and plotted as % cells with cilia. Downregulation of *trip6* reduced the number of cells with cilia (p>0.0001 Fisher’s exact test) and affected both the length and morphology. “Puncta” symbolizes extremely short cilia consisting of basal bodies with unextended axoneme marked by ARL13B. (**B**) Procedure as in A, except that a blocking peptide (or a control peptide) fused with mCherry was introduced into Z310 cells. The peptide inhibits by competition the formation of TRIP6 homodimers. This lead to decrease in the number of cells with cilia (p>0.0001 Fisher’s exact test) and the length of the cilia. Quantitation of the effect on cilia as in A. For details on the quantification please refer to the Materials & Methods section. Scale bars: 10μm (A and B).

In contrast to ciliogenesis, downregulation of *trip6* in Z310 cells by siRNA did not abolish either adherens junctions nor focal adhesions (**Figure S3B** and **C**). At focal adhesions, residual TRIP6 was still visible indicating a longer protein half-life. Most likely, TRIP6 is stabilized when incorporated into focal adhesions, remaining longer than at adherens junctions. Thus, in cultured cells as in the brain, absence of TRIP6 did not affect adhesion complexes (compare with **Figure 3 4A**).

TRIP6 carries three C-terminal LIM domains with different interaction specificities. In addition, the N-terminal half harbors other additional motifs for binding proteins. The ability to act as an assembly platform is further increased by two dimerization domains (Diefenbacher et al., 2014). To examine whether dimerization was required for the ciliogenesis role of TRIP6, we blocked dimerization by overexpressing a peptide competing with the dimerization domains (Diefenbacher et al., 2014) in Z310 cells. The resulting phenotype was similar to those findings obtained after downregulation by siRNA: the number and length of the cilia were significantly reduced upon transfection with the blocking peptide as compared to transfection with a control scrambled version of the peptide (**Figure 7B**). As in the downregulation of TRIP6 the block of dimerization caused “puncta” appearance of severely reduced cilia size, with similar quantification as in Figure 7A. These data suggest that TRIP6 requires the full complement of protein binding options, and that it may serve as an assembly factor for multiprotein complexes involved in cilium formation.

Similarly to the brain cilia phenotype (**Figure 5C**), super resolution microscopy of the cilia of Z310 cells revealed a punctate distribution of TRIP6 along the length of the ciliary axonema, somewhat differing in distribution between individual cilia (arrows in **Figure 8**, the examples of cilia shown represent a total number of 28 cilia with similarly differing TRIP6 distribution). This is suggestive of “snapshots” taken during transport. In contrast, the structural cilia component acetylated tubulin, as well as ARL13B (Roy et al., 2017), were more uniformly distributed along cilia. TRIP6 also co-localized with γ-tubulin at the basal bodies (**Figure 8B**). Compared to TRIP6, CLUAP1 was predominantly located at the distal and proximal ends of the cilia (**Figure 8C**), in agreement with previous reports (Botilde et al., 2013). Downregulation of *trip6* by siRNA caused shortening of the cilia (example shown in **Figure 8D** middle panels; a total of 20 and 15 cilia were imaged for siControl and siTrip6 respectively). Pericentrin, which marks pericentriolar material surrounding the basal bodies, was not affected. Because the downregulation of *trip6* was not quantitative due to incomplete transfection efficiency, cells with full-length cilia were detected in the same culture. These expressed TRIP6 along the axoneme and basal body (cilium example at bottom panel of **Figure 8D**) similar to siControl cells (**Figure 8D top panel**).

**Figure 8:**
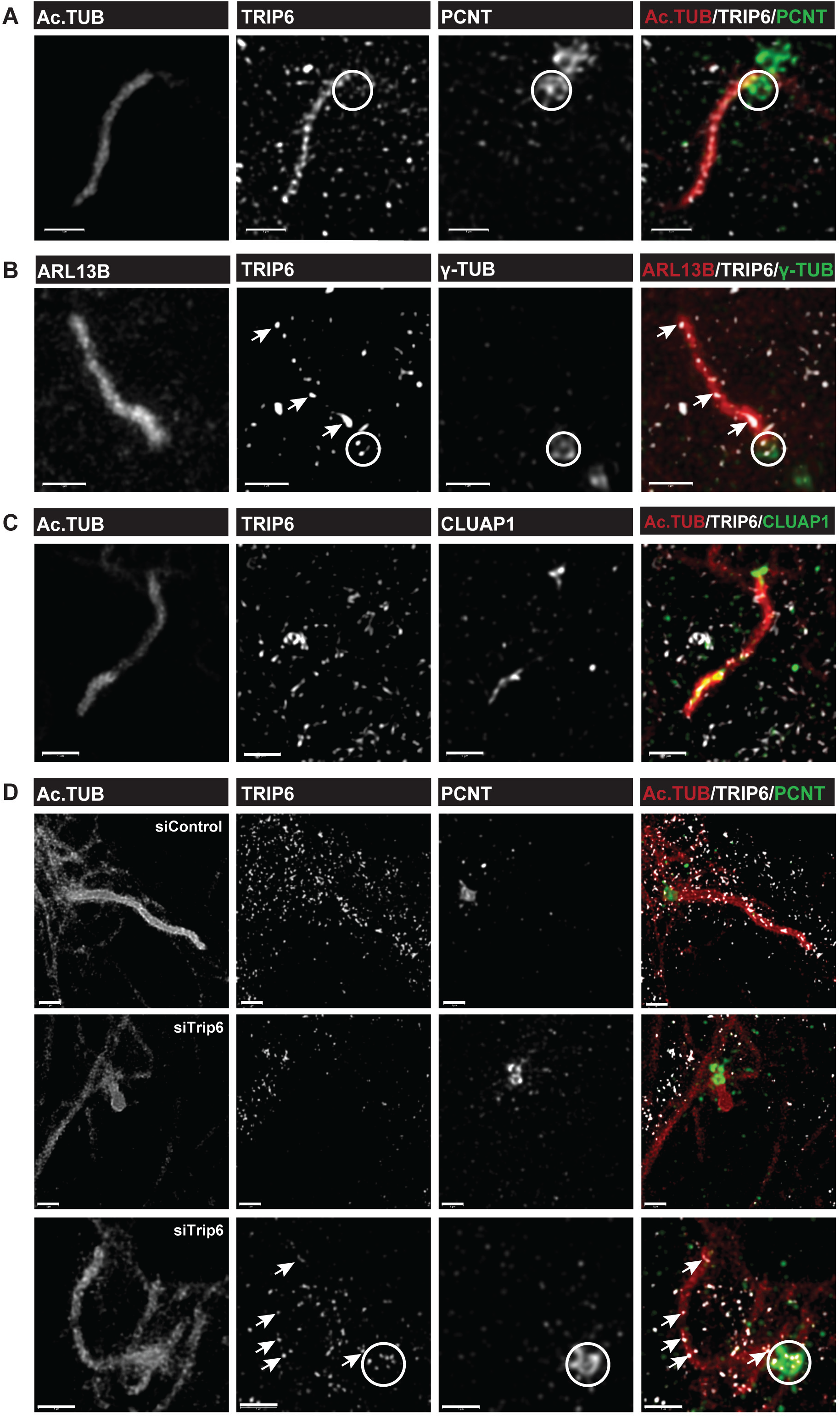
Super resolution imaging of choroid plexus epithelial cell line cilia. Cilia formation was induced in Z310 cells as in Figure 7. Different individual cilia were selected in **A** through **D**. Acetyl-tubulin (in **A**, **C** and **D**) and ARL13B (**B**) stained the axonemes uniformly, while TRIP6 was localized in dot-like association and in different pattern from one cilium to the next (examples indicated by arrow). The rings indicate the basal bodies, defined by localization of γ-TUBULIN (γ-TUB) and PERICENTRIN (PCNT) (**A** and **B**). Partial downregulation of *trip6* by siRNA (not the siControl **D**: upper panel) reduced the length of the axoneme (**D** middle panel) in those cells where TRIP6 was depleted from the cilium (**D**, middle panels). In cilia where TRIP6 continued to localize due to only partial downregulation by *trip6*siRNA (**D** bottom panel) the cilia elongated and localization of TRIP6 along the axoneme (arrows) and basal body (circle) was observed. Scale bars: 1μm (A, B, C and D).

## Discussion

In this work, we report that the LIM domain protein TRIP6 is required for proper differentiation of ependymal cells and choroid plexus in the developing mouse brain. Furthermore, we have uncovered a role for TRIP6 in ciliogenesis in the brain.

Surprisingly, in the *trip6* knockout animal, the ciliopathy-like phenotype was restricted to the brain, although TRIP6 is expressed almost ubiquitously in the organism both during early development and in the adult (UniProtKB Q9Z1Y4-TRIP6_MOUSE; https://www.proteinatlas.org/ENSG00000087077-TRIP6/tissue). Given their overlapping molecular properties and similarities in their structure, one may assume that in mice with *trip6* deletion, other members of the ZYXIN family of LIM domain proteins substitute for TRIP6 function in ciliogenesis. This hypothesis is supported by the observation that in Xenopus embryo the knockdown of WTIP, a ZYXIN family member, altered the location of basal bodies and reduced cilium length (Chu et al., 2016). A role for other ZYXIN family members in ciliogenesis is further plausible by the detection of ZYXIN, LPP as well as TRIP6 in the ciliary membrane-associated proteome of inner medullary collecting duct cells (IMCD3; Kohli et al., 2017). Our discovery of TRIP6 as an essential component of ciliogenesis was possible, because TRIP6 is the only member of the ZYXIN family of LIM domain proteins that is expressed during the differentiation of choroid plexus and ependymal cells (between E14.5 and birth, Figure 1 and mouse genome informatics atlas, Gray et al., 2004). Therefore, other family members cannot compensate for the loss of TRIP6 at this stage of brain development.

The differentiation and ciliogenesis defects in the developing brain of the trip6^−/−^mice are associated with severe hydrocephalus. However, hydrocephalus was observed in only 43% of the knockout animals, whereas the impaired differentiation of ependymal cells and of the choroid plexus, as well as the defective ciliogenesis were fully penetrant. These observations clearly suggest that hydrocephalus is not a direct consequence of the loss of TRIP6, but rather a secondary effect. Furthermore, the enlargement was restricted to the lateral ventricles, suggesting an occlusion of CSF passage downstream of these ventricles. This phenotype strikingly resembles human genetically determined non-communicating (obstructive) hydrocephalus, a condition caused by occlusion of CSF passage, which represents the most frequent hydrocephalus aetiology besides CSF overproduction (reviewed by Kousi & Katsanis, 2016). Thus, it seems that the obstructive hydrocephalus observed upon the loss of TRIP6 is caused by a cerebrospinal fluid (CSF) circulation block in the foramen of Monro (upstream of the third ventricle) and/or in the aqueduct of Sylvius, between the third and fourth ventricles. Given the reported role of TRIP6 in both focal adhesion and adherens junctions based on *in vitro* experiments (Guryanova et al., 2005; Takizawa et al., 2006; Bai et al., 2007), one logical hypothesis could have been that the loss of TRIP6 promoted a detachment of ependymal cells, ultimately occluding the aqueduct. Such a mechanism has indeed been reported in cases of congenital hydrocephalus (Wagner et al., 2003; Páez et al., 2007; Willems et al., 1987; Rosenthal et al., 1992) as well as in a number of animal models of defective cell adhesion (Yamamoto et al., 2013; Rakotomamonjy et al., 2017; Tissir et al., 2010; Guerra et al., 2015). In the trip6^−/−^mouse, the ependymal layer indeed shows patches of interruption compatible with ependymal cell detachment from the basement membrane (not shown). However, the loss of TRIP6 did not seem to hamper focal adhesion and adherens junctions. One possibility is that the defective differentiation combined with hampered ciliogenesis somehow reduced the adhesion strength of ependymal cells. This might in turn lead to delamination, clotting of the delaminated material and eventually hydrocephalus.

Upon deletion of *trip6*, we observed mal-differentiation of the ependyme and choroid plexus as well as ciliogenesis defects. However, it is not clear whether these phenotypes are linked, i.e. whether the differentiation defects are a consequence of the compromized cilia formation or are caused by *trip6* deletion independently. TRIP6 also acts as a transcription regulator via a nuclear isoform (Kassel et al., 2004; Diefenbacher et al., 2008; Diefenbacher et al., 2014; Kemler et al., 2016) and thus could regulate differentiation programmes. However, we have not detected nuclear TRIP6 in the brain cells examined. Nevertheless, for transcription regulatory functions minute levels may suffice for large effects. Expression below detection level might still be required for the differentiation of the ependyme. Furthermore, a transcriptional role for TRIP6 has been postulated in the regulation of proliferation, in particular in cancer (Willier et al., 2011, Miao et al., 2016; https://www.proteinatlas.org/ENSG00000087077-TRIP6/pathology). A putative role for TRIP6 in the regulation of proliferation has also been reported in the context of brain development: *in vitro*, postnatal neuronal progenitor cells suffered from poor proliferation upon knocking down TRIP6 (Lai et al., 2014). However, in the *trip6*^−/−^mouse embryo, we have not observed any obvious proliferation defects by gross examination and by Ki67 staining (not shown). Adult neuronal progenitors, although a minority, express several members of the ZYXIN family (Zywitza et al., 2018). Compensation for loss of TRIP6 could conceal a proliferation defect in the adult.

Our data point to a role for TRIP6 in the formation of both motile (*in vivo* analysis) and primary cilium (induction in cell culture). These two classes of cilia differ in their structure, development and function (reviewed by Narita & Takeda, 2015; Malicki & Johnson, 2017). While motile cilia primarily move extracellular fluid, e.g. CSF, the single so-called primary cilium is immotile and serves as a signalling centre (Goetz & Anderson, 2010). However, despite their functional differences, multiple structural studies have revealed their morphological similarities, and proteome analysis identified shared components (Narita et al., 2012). Thus, TRIP6 is apparently a shared component. A possible function of TRIP6 in the formation of both motile and primary cilium could be mediated through the cytoskeleton, given its described regulatory function in actin cytoskeleton dynamics (Sanz-Rodriguez et al., 2004; Guryanova et al., 2005; Bai et al., 2007). Interestingly, adhesion complexes such as those in focal adhesions are found in basal bodies, thus linking cilia with the actin cytoskeleton (Antoniades et al., 2014). Actin polymerization indeed affects primary cilium length and function (Kim et al., 2015; Drummond et al., 2018). Thus, a putative function of TRIP6 in ciliogenesis via a regulation of the actin cytoskeleton cannot be ruled out at this time. However, its distribution along the axonemes does not speak for an actin related mechanism.

Another shared feature of motile and primary cilia are the transport mechanisms into and out of the ciliary axoneme which are carried out by the same intraflagellar transport (IFT) protein complexes (reviewed by Reiter et al., 2012; Madhivanan & Aguilar, 2014). Thus, disrupting intraflagellar transport would affect both motile and primary cilia, as we observed upon *trip6* deletion. Our data indeed strongly suggest a role for TRIP6 in intraflagellar transport. TRIP6 partially co-localized with proteins involved in intraflagellar transport, namely CLUAP1/IFT38 and ARL13B. This is supported by previous observations that TRIP6 interacts with CLUAP1 in cytoplasmic complexes of HEK293T cells (Beyer et al., 2018). and with additional centrosome and cilium-transition zone related proteins (Gupta et al., 2015; http://prohits-web.lunenfeld.ca). Furthermore, the punctate pattern of TRIP6 localization in individual cilia differed from one cilium to the other, as if TRIP6 containing transport complexes were fixed at different stages of the dynamic ciliogenic process, both *in vivo* and after induction by serum deprivation in cell culture. Interestingly, our results show that TRIP6 needs to homodimerize for proper function in ciliogenesis. Dimerization, which doubles the protein interaction sites, most likely allows TRIP6 to interact simultaneously with several cargo proteins in addition to CLUAP1/IFT38 and ARL13B. TRIP6 was identified to be in the proximity of a variety of proteins that localize to the satellite and appendages of the centrioles and within the centrioles themselves along with proteins from the transition zone of the primary cilium – this wide distribution of interactors is a characteristic of an adaptor protein/assembly factor, supporting our conclusions (Gupta et al., 2015; http://prohits-web.lunenfeld.ca). Conversely, our results show that TRIP6 is not required for the normal localization of γ-tubulin and pericentrin which marks the pericentriolar material surrounding the basal bodies, indicating that the regulatory stimuli inducing cilium formation are not altered. Thus, given that active transport along the ciliary axoneme is required for cilium formation (Davenport et al., 2007; Banizs et al., 2005; Town et al., 2008; Botilde et al., 2013), the ciliogenesis defects (reduced cilia number and size) observed in the absence of TRIP6 are most likely related to reduced intraflagellar transport.

In conclusion, this work has unveiled a novel role for TRIP6 in ciliogenesis, which is critical for brain development. This expands the spectrum of functions of this LIM domain protein beyond the regulation of adhesion, migration and proliferation. That a protein assembly factor is required for ciliogenesis, is a novel discovery.

## Materials and Methods

### Generation of trip6 knockout mice

Mice carrying trip6 deleted alleles were generated by ES cell gene targeting as depicted in Figure S1. Briefly, E14.1 mouse embryonic stem (ES) cells were electroporated with the linearized targeting vector and correct targeting events were identified by Southern-blot hybridization. Selected ES cell clones were subsequently electroporated with the pMC-Cre expression plasmid for deletion of the floxed neo-resistance cassette. Finally, blastocyst microinjection and transfer into foster mice was employed to generate knockout mouse lines. The resulting trip6 knockout mouse line was backcrossed to C57BL/6J mice for ten generations. All animal experiments were conducted according to German animal welfare legislation.

### Mouse colony maintenance and genotyping

The trip6 knockout colony was maintained by breeding heterozygous animals. To obtain homogeneous lines, routine backcrossing to the parental C57BL/6J strain was employed. Animals were provided with standard laboratory chow and tap water ad libitum and kept in accordance with local regulations (TLLV Thüringen, Erfurt, Germany) at constant temperature (22°C) and light cycle (12hr-light, 12hr-dark). Mouse colony maintenance and breeding were performed in accordance with regulations of the relevant authority (TLV; Thüringen, Germany) and under the oversight of the FLI Animal Welfare Committee. Postnatal day 4 (P4) mice were sacrificed by decapitation whereas mice older than ten days were sacrificed by CO_2_ inhalation. The genotype of mice was determined by PCR on DNA extracted from tail tissue using the following primers- trip6-28F: TCA CCT TTT CTC CCT TGC CTG CCT, trip6-29R: GGT ACC CCC GGA GGC TGA TAA CAG, trip6-30R: GCT TAT CGA TAC CGT CGA CCT CGA

### mRNA expression in mouse brain sections by in-situ hybridization

A cDNA fragment corresponding to nt995-1537 of mouse trip6 cDNA (GenBank accession number NM_011639.3) was generated by PCR and subcloned into the pGEM-T Easy Vector (Promega, Madison, WI). Radiolabeled riboprobes were generated by using [35S]UTP (Hartmann Analytik, Braunschweig, Germany) as substrate for the in vitro transcription reaction. ISH was carried out as published elsewhere (Heuer et al., 2000). Briefly, frozen, 20 μm thick frontal brain sections were fixed with 4% phosphate-buffered paraformaldehyde solution (pH 7.4) at room temperature (RT) for 60 minutes and rinsed with PBS (pH 7.4). Tissue sections were treated with 0.4% Triton-PBS for 10 minutes (RT) and then incubated in 0.1M triethanolamine, pH 8, containing 0.25% v/v acetic anhydride for 10 minutes. Following acetylation, sections were rinsed several times with PBS, dehydrated by successive washings with increasing ethanol concentrations, and air-dried. Radioactive 35S-labeled riboprobes were diluted in hybridization buffer (50% formamide, 10% dextran sulfate, 0.6 M NaCl, 10 mM Tris/HCl pH 7.4, 1 3 Denhardt’s solution, 100 mg/ml sonicated salmon-sperm DNA, 1 mM EDTA-di-Na, and 10 mM dithiothreitol) to a final concentration of 25000 cpm/μl. After application of the hybridization mix, sections were cover-slipped and incubated in a humid chamber at 58°C for 16 hours. Following hybridization, coverslips were removed in 2× standard saline citrate (SSC; 0.3 M NaCl, 0.03 M sodium citrate, pH 7.0). The sections were then treated with RNase A (20 mg/ml) and RNase T1 (1 U/ml) at 37°C for 30 minutes. Successive washes followed at RT in 1×, 0.5×, and 0.2× SSC for 20 minutes each and in 0.2× SSC at 60°C for 1 hour. The tissue was dehydrated and, for microscopic analysis, sections were dipped in KodakNTB nuclear emulsion (Kodak, Rochester, NY) and stored at 4 °C for 32 days. Autoradiograms were developed in Kodak D19 for 5 minutes, fixed in Rapid Fix (Kodak) for 10 minutes and analyzed under dark-field illumination. Experiments carried out using the respective sense probe did not produce any hybridization signals.

### Quantification of percentage of mice with hydrocephalus

The analysis was performed on an age-matched cohort, the details described here are - genotype: n^H^= number of mice with hydrocephalus; n=total mice analysed, number of females, number of males, age range of the cohort.

**1**) trip6^+/+^: n^H^=0; n=162, ♀=85, ♂=77, 5-417 days; **2**) trip6^+/−^: n^H^=0; n=186, ♀=94, ♂=92, 3-460 days;

**3**) trip6^−/−^: n^H^=104; n=240, ♀=125, ♂=115, 3-425 days. Hydrocephalus was scored on the basis of gross examination, an example of which has been shown in Figure 2A.

### MRI

To image ventricle size and morphology in early post-natal mice with normal neurogenesis and under pathological conditions in situ, a high resolution (150μm isotropic) MRI protocol was applied. Scans were performed on a clinical 3T scanner (Magnetom Trio, Siemens Healthcare), using a dedicated mouse head coil. A Siemens SPACE sequence with a constant flip angle was employed to acquire images with a resolution of 0.2 mm × 0.2 mm × 0.16 mm using the following parameters: echo time TE = 125 msec, repetition time TR = 1900 msec, bandwidth = 130 Hz/px, Turbo Factor TF = 65 and integrated fat saturation. MRI data were processed using the software syngo fastView. For depiction of multicolored axial images multiplanar reconstructions from the original 3D MRI data set were prepared.

### Histological processing and Immunofluorescence labelling

**Paraffin embedding** of mouse brain was performed after fixation in 4% phosphate-buffered paraformaldehyde solution (pH 7.4) for 48 hours at 4°C. For sagittal sectioning, the brain was cut in two halves along the midline before embedding in paraffin, whereas for coronal sectioning the brain was embedded intact in paraffin.

**Haematoxylin & Eosin staining** was performed according to standard protocol on 5μm-thick brain sections, prepared from the paraffin-embedded tissue. Images were acquired at a VS110 microscope (Olympus) with a 20× objective.

**Immunofluorescence labelling** was performed according to Li et al., 2015. Briefly, 5 μm-thick paraffin-embedded brain sections from P4 mice were deparaffinized and rehydrated (trip6^+/−^: n=5 and trip6^−/−^: n=5). Heat induced antigen retrieval was performed in citrate buffer (10mM, pH 6.0) in a decloacking chamber (Biocare Medical) by preheating at 95°C for 10 minutes followed by 125°C for 25 minutes. After cooling the sections were washed in 1x phosphate buffered saline (PBS, pH7.4), incubated with immunohistochemistry blocking buffer (5% BSA, 5% goat serum, 0.1% triton X-100) for 1 hour followed by primary antibody (Table S1) incubation at 4°C overnight. The samples were washed in PBS and incubated with secondary antibodies (Table S2) for 1 hour. The nuclei were counter-stained with 1 μg/ml DAPI (4′,6-diamidino-2-phenylindole, Sigma) for 5 minutes. After washing the sections were cover-slipped in Mowiol® 4-88 (Sigma) mounting medium containing 1 mg/ml p-phenylenediamine (Sigma).

**Imaging** was performed on an Axio Imager microscope (Carl Zeiss) equipped with ApoTome-slider and a 12-bit grayscale cooled CCD AxioCamMRm camera. Z-stacks were acquired and the representative images shown in the figures are maximum intensity projections generated using ZEN software (Carl Zeiss). Representative images were brought to a resolution of 300 ppi using Adobe Photoshop (Adobe) and the area of interest was cropped.

### Cell culture and induction of cilia

Immortalized murine choroid plexus cell line Z310 (RRID:CVCL_F753) were kindly provided by Professor Zheng, Purdue University. Z310 cells were cultured in DMEM high glucose (4.5 g/l) supplemented with 10% FBS and 2 mM glutamine at 37°C in 5% CO_2_. As described, the cell line had been characterized by genotyping, morphology, growth assays and expression of the choroidal epithelium-specific marker TTR (Zheng & Zhao, 2002).

**Ciliogenesis** was induced by starvation at 37°C in 5% CO_2_ for 24 hours. For starvation, the growth medium was changed to DMEM low glucose (1 g/l) without FBS and supplemented with 2 mM glutamine.

### Transfections

**siRNA transfection** was performed using ON-TARGETplus siRNA targeting the Human trip6 ORF (Dharmacon). siRNA sequences are listed in table S3. Pools of siRNAs duplexes were employed, with each pool containing 2 independent oligonucleotides targeting trip6. A pool of non-targeting siRNAs was used as negative control. Lipofectamine RNAiMax (Thermo Fisher) was used for transfection according to the manufacturer’s protocols. First, Z310 cells were reverse-transfected with the siRNA pool, subsequently at 48 hours, forward transfection with siRNA and induction of ciliogenesis (via starvation) were performed simultaneously.

**Plasmid transfection** was performed with Lipofectamine2000 (Thermo Fisher) according to manufacturer’s protocol. The pcDNA3.1 constructs coding for the dimerization inhibitory peptide (amino acid 253-265 of human TRIP6 which is conserved in mouse and rat) and the scrambled control peptide tagged with mcherry have been described previously (Diefenbacher et al., 2014). Briefly, Z310 cells were first transfected with the relevant constructs and 48 hours later the cells were starved for 24 hours to induce ciliogenesis. The cells were subsequently fixed and permeabilized for immunofluorescence labelling

### Immunofluorescence labelling

Z310 cells were labelled as described previously (Li et al., 2016). Briefly, both transfected and untransfected cells were grown on glass coverslips and fixed with 3% paraformaldehyde and permeabilized with ice-cold methanol. Subsequent to treatment with NaBH_4_ (0.01% in PBS), washing in PBS, the coverslips were incubated with blocking buffer (1% BSA, 2% FCS in PBS) for 30 minutes, followed by primary and secondary antibodies for 45 minutes each. The nuclei were stained with DAPI (1mg/ml for 5 minutes) and the coverslips were mounted on glass slides in Mowiol® 4-88 (Sigma) mounting medium containing 1mg/ml p-phenylenediamine (Sigma).

### Image acquisition and quantification

Images were acquired on a Zeiss Axio Imager microscope (Carl Zeiss) equipped with ApoTome-slider, an 63×1.4 NA objective and and a 12-bit grayscale cooled CCD AxioCamMRm camera. The cilia were imaged as z-stacks encompassing the z positions from the basal bodies up to the tip of the axoneme.

Quantification of number of cells with cilia was performed on Maximum Intensity Projections (MIP) for each image using ZEN software (Carl Zeiss). The ‘Grid’ and ‘Events’ functions from the ZEN (desk) software were used to facilitate the counting of the total number of nuclei and the number of cilia in each field. A cilium was counted only when the basal body (gamma-tubulin) and the axoneme (acetyl-tubulin or ARL13B) were visible in order to avoid false signals (common for acetyl-tubulin). The nuclei touching the periphery of the field were excluded unless a cilium emerged from the same cell. The percentage of cells with cilia were calculated from three replicate experiments. A minimum of 7 fields were counted for each condition (controlsiRNA vs trip6siRNA and Control vs Blocking peptide) in each replicate experiment. Cilia length was measured using the ‘spline curve’ measurement tool of the ZEN software. The mean and standard deviations were calculated from n=59 cells for controlsiRNA and n=75 cells for trip6siRNA (Figure 7A), and n= 62 and n=56 for blocking peptide experiment (Figure 7).

### Super-resolution microscopy

Super-resolution imaging was performed on a Leica TCS SP8 X-White Light Laser confocal microscope equipped with an inverted microscope (DMI 8, Leica), a 100× objective (HC PL APO CS2 100×1.4 oil), Hybrid detectors and HyVolution-II software. Images were acquired as suggested by the HyVolution module within the LAS microscope software (Leica). Pre-settings in the HyVolution mode were established to achieve the highest possible optical resolution. The super-resolved confocal stacks were deconvolved in Scientific Volume Imaging (SVI)/Huygens deconvolution professional suite software with GPU acceleration.

HyVolution is based on a combination of confocal imaging using sub-Airy pinhole sizes (0.45 - 0.6 Airy Units, depending on the fluorophores used) with subsequent computational image deconvolution as initially described by Schrader et al., (1996) and Lam et al., (2017). With HyVolution, a lateral resolution of 140 nm can be achieved, which is approximately 1.6 times higher than the resolution achievable with conventional confocal microscopy. For HyVolution imaging, IF labelling was performed as described previously for Z310 cell and Paraffin embedded mouse brain tissues. TRIP6 was labelled along with basal body markers: γ-TUBULIN and PERICENTRIN, and cilia markers: Acetylated αTUBULIN, ARL13B and CLUAP1 (antibody details in Table S1). The secondary antibodies used are listed in Table S2. The samples were mounted in ProLong Gold antifade mountant (Thermo Fisher). HyVolution images were acquired using the 488 nm, 546 nm and 647 nm laser lines of the white-light laser to excite Alexa Fluor 488, Alexa Fluor 546 and Alexa Fluor 647 dyes, respectively. Fluorescence emission was recorded on a hybrid detector at 503 nm – 547 nm, 558 nm – 604 nm, and 654 nm – 719 nm, respectively, employing sequential scan imaging to avoid bleed-trough and/or cross-talk between spectral channels. DAPI was excited using a 405 nm laser and detected using a PMT-detector (430 nm – 502 nm).

### Statistical analysis

Statistical significance of number of mice with hydrocephalus was analysed by the chi-squared test. The statistical significance of the difference in the number of cells with cilia in controlsiRNA vs trip6siRNA and Control vs Blocking peptide was tested by Fisher’s exact test. The two tail Student’s t-test was used to analyse the difference in the length of the cilia between controlsiRNA vs trip6siRNA and Control vs Blocking peptide.

---

**Table S1.** Details of primary antibodies used for immunofluorescence labelling.

|  | Primary antibody | Supplier (Cat#) | Dilution |
| --- | --- | --- | --- |
| 1 | Anti-Trip6 (A15) | Santa cruz biotechnologies (sc-34976) | 1:50 |
| 2 | Anti-Acetylated Tubulin | Sigma (T6793) | 1:100 |
| 3 | Anti-gamma tubulin (GTU88) | Sigma (T6557) | 1:200 |
| 4 | Anti- Paxillin | Transd. Lab (P13520) | 1:1000 |
| 5 | Anti-β-Catenin | Sigma (C2206) | 1:500 |
| 6 | Anti-ARL13B | Proteintech (17/11-1-AP) | 1:100 |
| 7 | Anti-CLUAP1 | Proteintech (17470-1-AP) | 1:100 |
| 8 | Anti-β-Catenin (12F7) | Santa cruz Biotechnology (sc-59737) | 1:100 |
| 9 | Anti-S100β | DAKO (GA504) | 1:100 |
| 10 | Anti-Pericentrin | Biolegend (923701) | 1:100 |

**Table S2.** Details of secondary antibodies used for immunofluorescence labelling.

|  | Secondary antibody | Supplier (Cat#) | Dilution |
| --- | --- | --- | --- |
| 1 | Donkey anti-Mouse IgG (H+L) Highly Cross-Adsorbed Secondary Antibody, Alexa Fluor 546 | Life technologies (A10036) | 1:100 |
| 2 | Donkey anti-Goat IgG (H+L) Cross-Adsorbed Secondary Antibody, Alexa Fluor 546 | Life technologies (A11056) | 1:100 |
| 3 | Donkey anti-Rabbit IgG (H+L) Highly Cross-Adsorbed Secondary Antibody, Alexa Fluor 546 | Life technologies (A10040) | 1:100 |
| 4 | Donkey anti-Mouse IgG (H+L) Highly Cross-Adsorbed Secondary Antibody, Alexa Fluor 488 | Life technologies (A21202) | 1:100 |
| 5 | Donkey anti-Goat IgG (H+L) Cross-Adsorbed Secondary Antibody, Alexa Fluor 488 | Life technologies (A11055) | 1:100 |
| 6 | Donkey anti-Rabbit IgG (H+L) Highly Cross-Adsorbed Secondary Antibody, Alexa Fluor 488 | Life technologies (A21206) | 1:100 |
| 7 | Donkey anti-Rabbit IgG (H+L) Highly Cross-Adsorbed Secondary Antibody, Alexa Fluor 647 | Life technologies (A31573) | 1:100 |
| 8 | Donkey anti-Mouse IgG (H+L) Highly Cross-Adsorbed Secondary Antibody, Alexa Fluor 647 | Life technologies (A31571) | 1:100 |

**Table S3.**
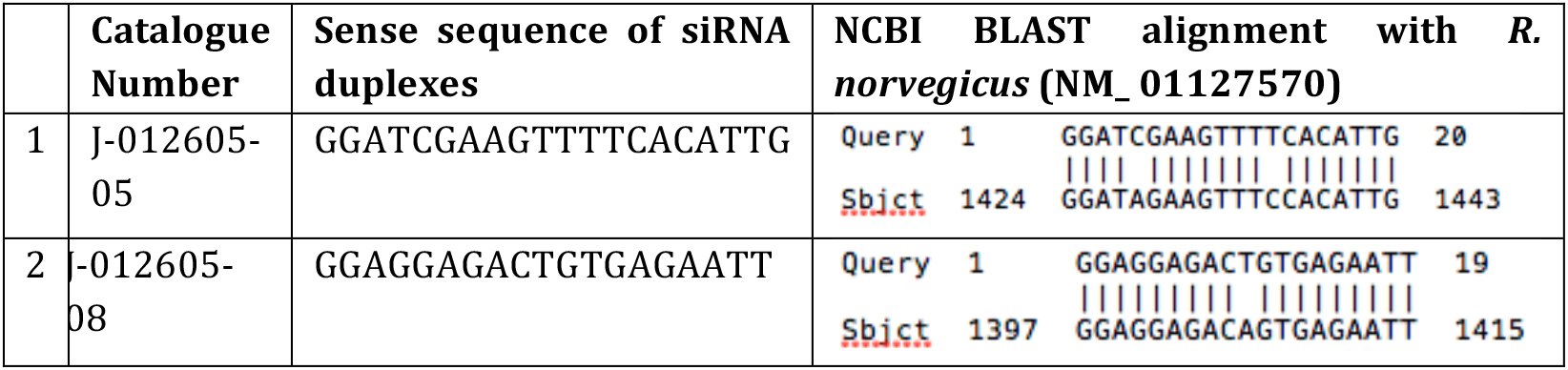
Sequence of siRNA targeting *trip6*.

## Legends

Figure S1: **Generation of** *trip6* **deleted mouse.** (**A**) *trip6* gene targeting strategy. (**B**)-(**C**) Southern blot analysis of *trip6* targeted (**B**), trip6 deleted and trip6 floxed (**C**) mice.

Figure S2: **ISH controls and TRIP6 expression in E15.5 brain.** (**A**) *trip6* mRNA expression determined by ISH. Embryonic sagittal sections through total skull from *trip6*^+/+^and mice *trip6*^−/−^were hybridized with *trip6* sense and antisense probes. Trip6 antisense serves as proof of trip6 deletion, *trip6*^+/+^ sense as negative control of the ISH. (**B**) Postnatal coronal sections, ISH as in **A**. The mice in B are different from those in Figure 1, thus further confirming the ISH control data. (**C**) Immunofluorescence of VZ cells and choroid epithelium showing co-staining of cellular membranes with β-catenin, N-cadherin and Trip6. Oblique sections through the VZ show several adjacent layers of VZ cells.

Figure S3: **No effect of the** *trip6* **deletion on focal adhesions and adherens junctions.** The choroid plexus cell line Z310 was transfected with *trip6* siRNA or control siRNA and grown in culture to semi-confluence. (**A**) The success of *trip6* downregulation was determined by immunoblot. Two exposures show the partial (approximately 50%) loss of TRIP6 protein. The Ponceau stain served as loading control. (**B**) Colocalization of Trip6 with paxillin at focal adhesions in starved (induction of ciliogenesis) Z310 cells. Downregulation of *trip6* did not affect the focal adhesions. (**C**) Immunofluorescence staining of β-catenin. No loss of adherens junctions upon downregulation of *trip6* was seen during induction of ciliogenesis. Note that visualization of adherens junctions requires relatively high cell density, while the formation of focal adhesions are better visible at lower cell density. Scale bars: 10μm (B andC).

## Acknowledgements

The support of the Animal House, Histology and Imaging Core Facilities of the FLI is gratefully acknowledged. We are grateful to the members of our laboratories for various discussions, in particular Torsten Kroll, Huaibiao Li, Yu-Chieh Lin, Markus Winter, Marcel König and Ali Adnan. We thank Silke Schulz for genotyping and Ines Krumbein for MRI advice. We appreciated the help of Karl-Heinz Herrmann in the MRT analysis. The project has received funding from the Jung Stiftung für Wissenschaft und Forschung (Herrlich laboratory). R.H. received funding by the VELUX Stiftung (Switzerland). The FLI is a member of the Leibniz Association and is financially supported by the Federal Government of Germany and the State of Thuringia. The authors declare no competing financial interests.

## Author Contributions

PU generated the *trip6^−/−^* mouse; SN performed ISH; WKM collected the data and SS performed analysis for hydrocephalus quantification; RH & WKM performed MRI; LF performed histological analysis; SS and AT analyzed brain sections and IF; SS performed IF staining and imaging; SS performed *in vitro* experiments; SS & SM performed super resolution microscopy; PH, SS and AP designed the study; PG, ZQW & PH acquired animal licenses; PH, SS, OK, AP & PU wrote the manuscript; PH, AP, ZQW, HH & OK supervised the study; all authors edited the manuscript.

