## Supplementary figures and images for "Novel function of TRIP6, in brain ciliogenesis"

### Supplementary Figure 1

FigureS1 - Trip6 gene targeting

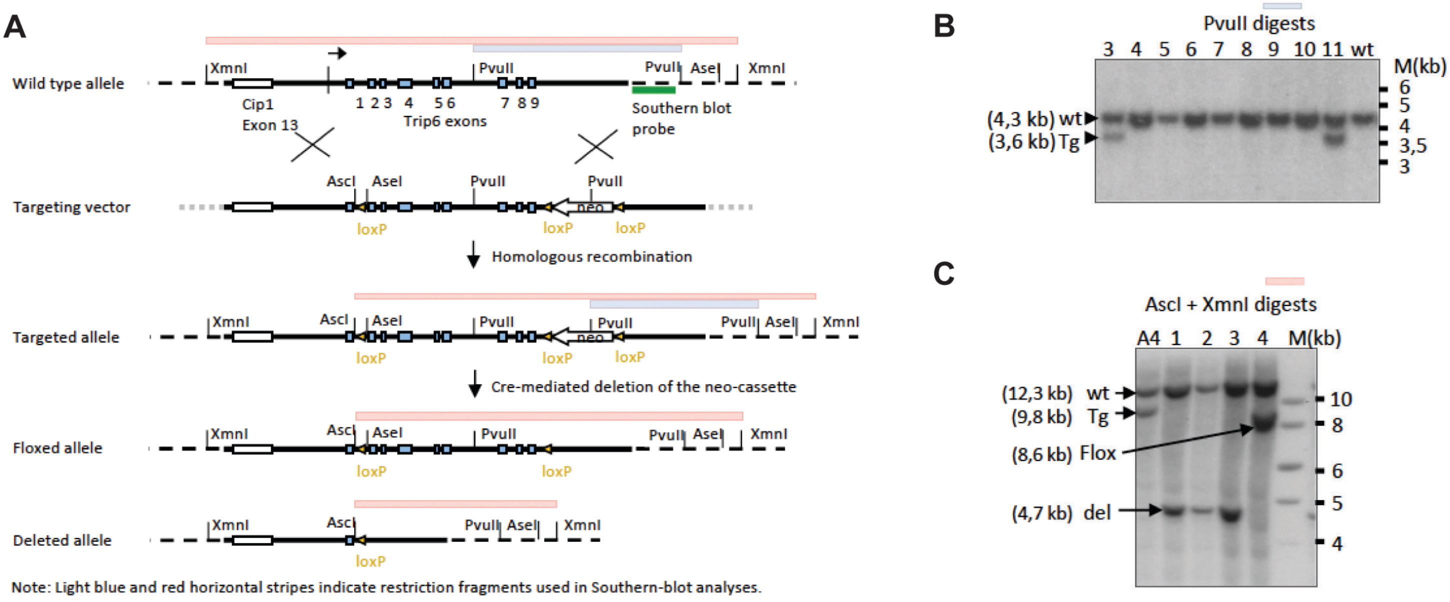

### Supplementary Figure 2

Figure S2 - ISH controls and Trip6 expression in E15.5 brain.

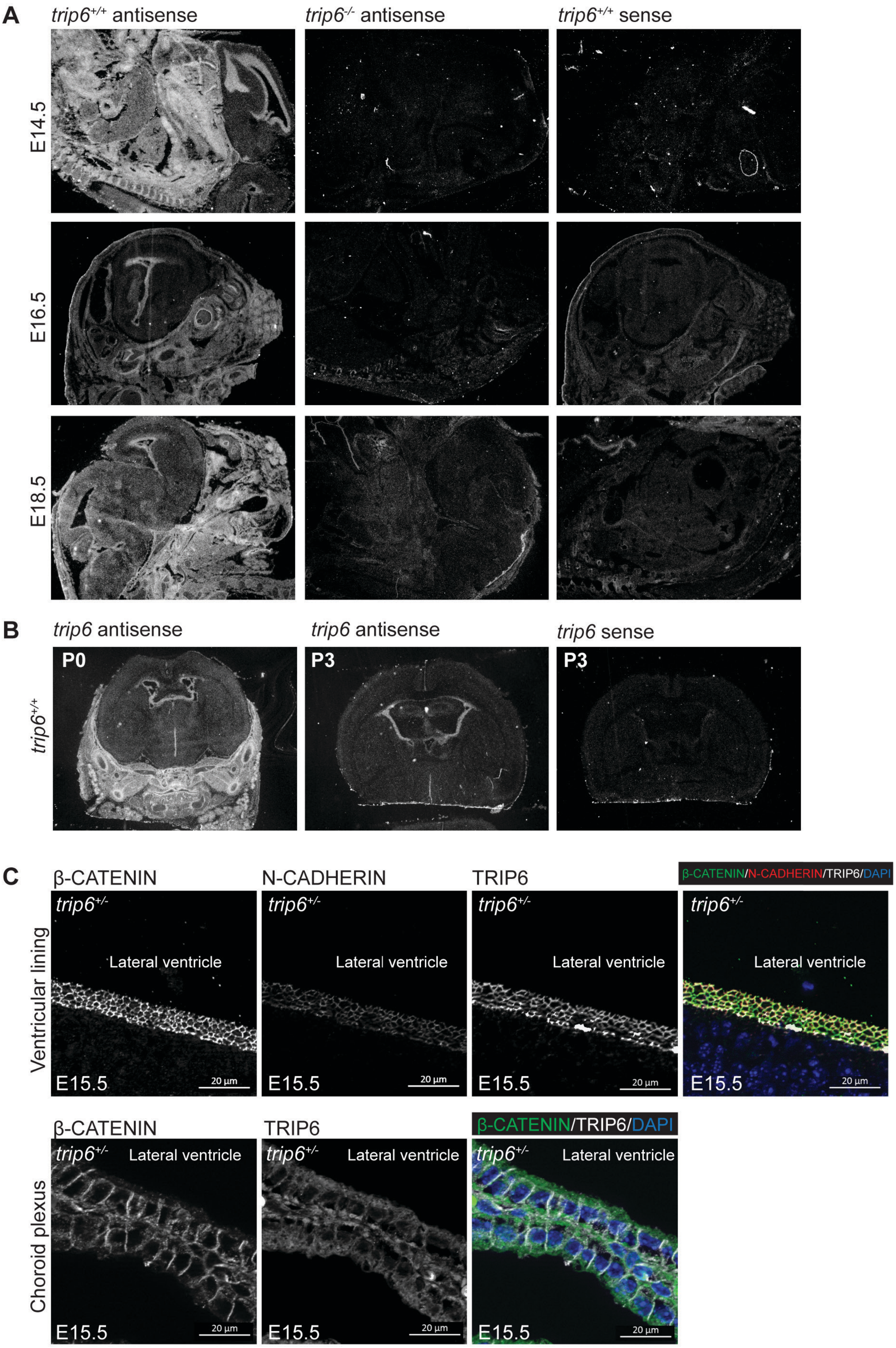

### Supplementary Figure 3

Figure S3 - No effect of the trip6 deletion on focal adhesions and adherens junctions.

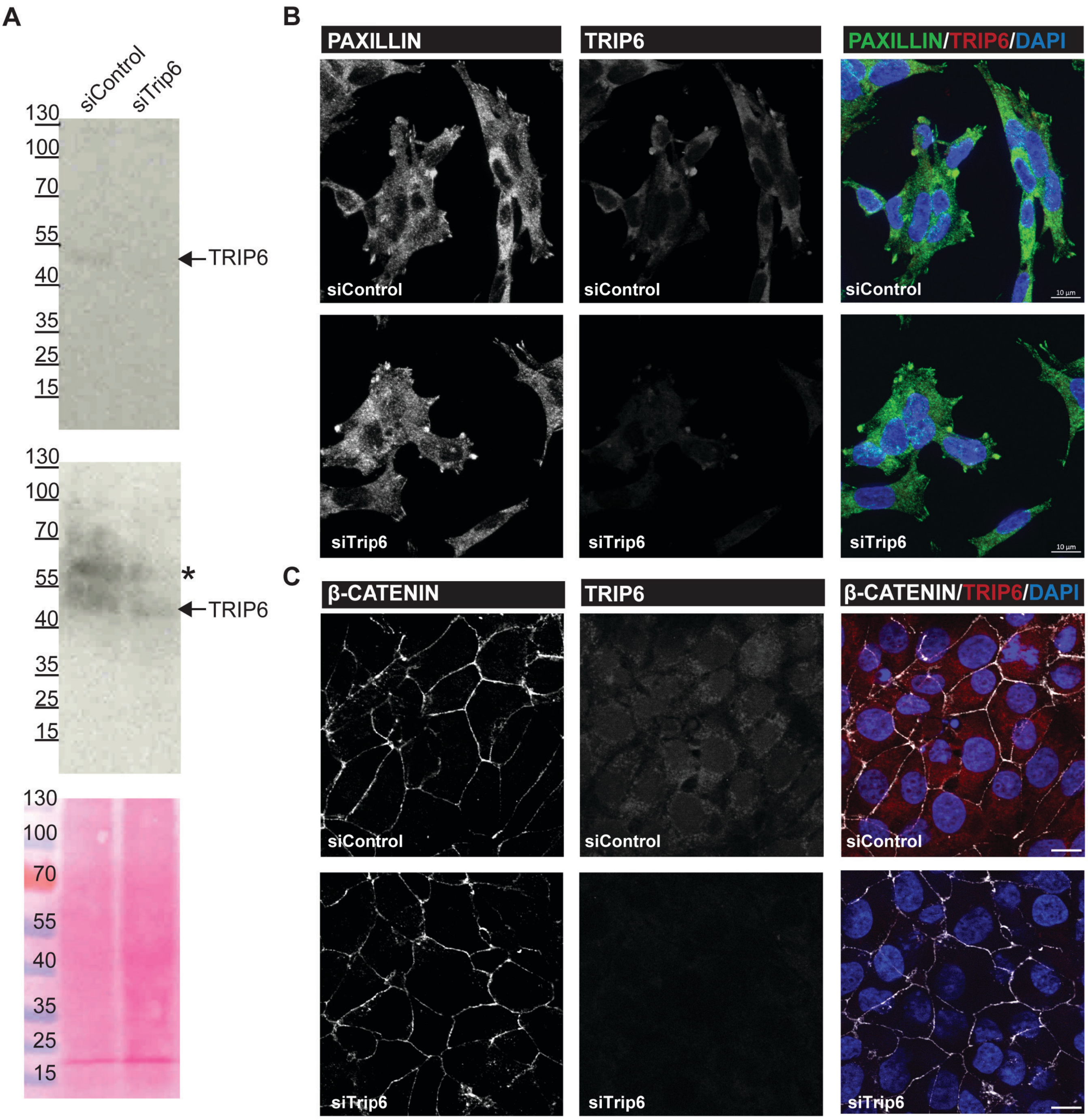
